# Cell type-specific adaptation of the SARS-CoV-2 spike

**DOI:** 10.1101/2023.12.20.572504

**Authors:** Marc Carrascosa-Sàez, María-Carmen Marqués, The IBV-Covid19-Pipeline, Ron Geller, Santiago F. Elena, Amal Rahmeh, Jérémy Dufloo, Rafael Sanjuán

## Abstract

SARS-CoV-2 can infect various human tissues and cell types, principally via interaction with its cognate receptor ACE2. However, how the virus evolves in different cellular environments is poorly understood. Here, we used experimental evolution to study the adaptation of the SARS-CoV-2 spike to four human cell lines expressing different levels of key entry factors. After 20 passages, cell type-specific phenotypic changes were observed. Selected spike mutations were identified and functionally characterized in terms of entry efficiency, ACE2 affinity, spike processing, TMPRSS2 usage, entry pathway and syncytia formation. We found that the effects of these mutations varied across cell types. Interestingly, two spike mutations (L48S and A372T) that emerged in cells expressing low ACE2 levels increased receptor affinity, syncytia induction, and entry efficiency under low-ACE2 conditions. Our results demonstrate specific adaptation of the SARS-CoV-2 spike to different cell types and have implications for understanding SARS-CoV-2 tissue tropism and evolution.

## Introduction

Severe acute respiratory syndrome coronavirus 2 (SARS-CoV-2) was identified in a pneumonia outbreak in Wuhan (China) in 2019^1,2^. Since then, it has circulated worldwide, causing millions of infections and deaths. The SARS-CoV-2 spike (S) glycoprotein is responsible for receptor binding, cell tropism and viral entry. It is composed of two subunits, S1 and S2, that are held together by non-covalent interactions in the mature virion. S1 is responsible for receptor binding and S2 mediates membrane fusion^3^. Spike activation is a multi-step process mediated by several cellular proteases. During viral egress, the spike precursor (S0) is first cleaved by furin at the S1/S2 junction, which contains a multibasic furin cleavage site (FCS)^4^. Further processing occurs at the S2’ site in target cells, exposing the fusion peptide (FP) and allowing viral entry. Cleavage at the S2’ site by the transmembrane protease serine 2 (TMPRSS2) triggers direct fusion with the plasma membrane^5^. Alternatively, SARS-CoV-2 can enter cells via clathrin-mediated endocytosis, which allows spike activation by cathepsins and fusion with the endosome membrane^6^. In addition to allowing fusion of cellular and viral membranes, spike-ACE2 interaction can induce cell-cell fusion, which has been suggested to play a role in viral spread, immune escape, and pathogenesis^7^.

The major SARS-CoV-2 receptor is human angiotensin-converting enzyme 2 (ACE2)^5^, which is expressed in various tissues, including the gastrointestinal tract, kidney, heart, respiratory tract, and female reproductive tract^8^. Accordingly, SARS-CoV-2 has a relatively broad tropism in cell cultures^9,10^ and *in vivo*^11^. Viral antigens have been detected in several tissues from COVID-19 patients, including trachea, lung, liver^12^, intestine^13^, kidney^14^, and central nervous system^15^. In addition to ACE2, other SARS-CoV-2 entry factors have been described, such as DC-SIGN/L-SIGN^16,17^, CD147^18^, neuropilin-1 (NRP1)^19,20^, and TMEM106B^21^, which may further extend viral tropism. However, to what extent this intra-host tropism determines the evolution of the SARS-CoV-2 spike is not well characterised.

Since its appearance in 2019, many SARS-CoV-2 linages have emerged and some have been designated as variants of concern (VOC), all of which harboured mutations in the S protein. Some of these changes have been associated to enhanced replication^22,23^, increased affinity for ACE2^24,25^, altered spike cleavage^26^, usage of alternative entry pathways^27^, or escape from neutralizing antibodies^28,29^. For example, the Omicron variant has shifted from TMPRSS2-dependent entry in previous strains to preferential entry via the endosomal pathway^30^. This has been suggested to change the tissue tropism of Omicron from the lower to the upper respiratory tract, thereby decreasing pathogenicity but increasing viral transmissibility. Such a change in tropism exposes the virus to new cellular environments that may alter its evolution.

Here, we sought to compare the adaptation of the SARS-CoV-2 spike to human cell lines expressing different levels of host entry factors. We used a recombinant, replication-competent vesicular stomatitis virus (VSV) system to experimentally evolve the S protein in four human cell lines, and identified adaptive mutations throughout the S protein. We characterized individual mutations in terms of entry efficiency, spike processing, activation by TMPRSS2, use of alternative entry routes and cell-cell fusogenicity. Finally, we showed that the effect of these substitutions on both cell entry and viral fitness is cell type-dependent.

## Results

### Adaptation of the SARS-CoV-2 spike to different human cell types

We produced a recombinant VSV in which the VSV glycoprotein was substituted with the SARS-CoV-2 spike (Wuhan-Hu-1 strain). The use of recombinant VSV has been shown to be a useful and relevant surrogate system to study many BSL3 and 4 viruses^31–33^, and particularly SARS-CoV-2 spike-dependent phenotypes like cell entry^34^, neutralization^34–36^, and adaptation^37,38^. Moreover, using spike-expressing recombinant virus for long-term passaging experiments in human cells allows to avoid using real virus and the potential emergence of gain-of-function mutations. We evolved this virus in four human cell lines: A549-ACE2 (A549-A), A549-ACE2-TMPRSS2 (A549-AT), Huh-7, and IGROV-1 (**Figure 1A**). These are well-characterized cell lines^39–43^ originating from different tissues and show large differences in the mRNA expression levels of key factors related to spike function such as ACE2, TMPRSS2, furin, and cathepsin L, as determined by RT-qPCR (**Figure 1B**). For instance, ACE2 mRNA levels spanned three orders of magnitude, with Huh-7 expressing the lowest and A549-AT the highest values.

**Figure 1.**
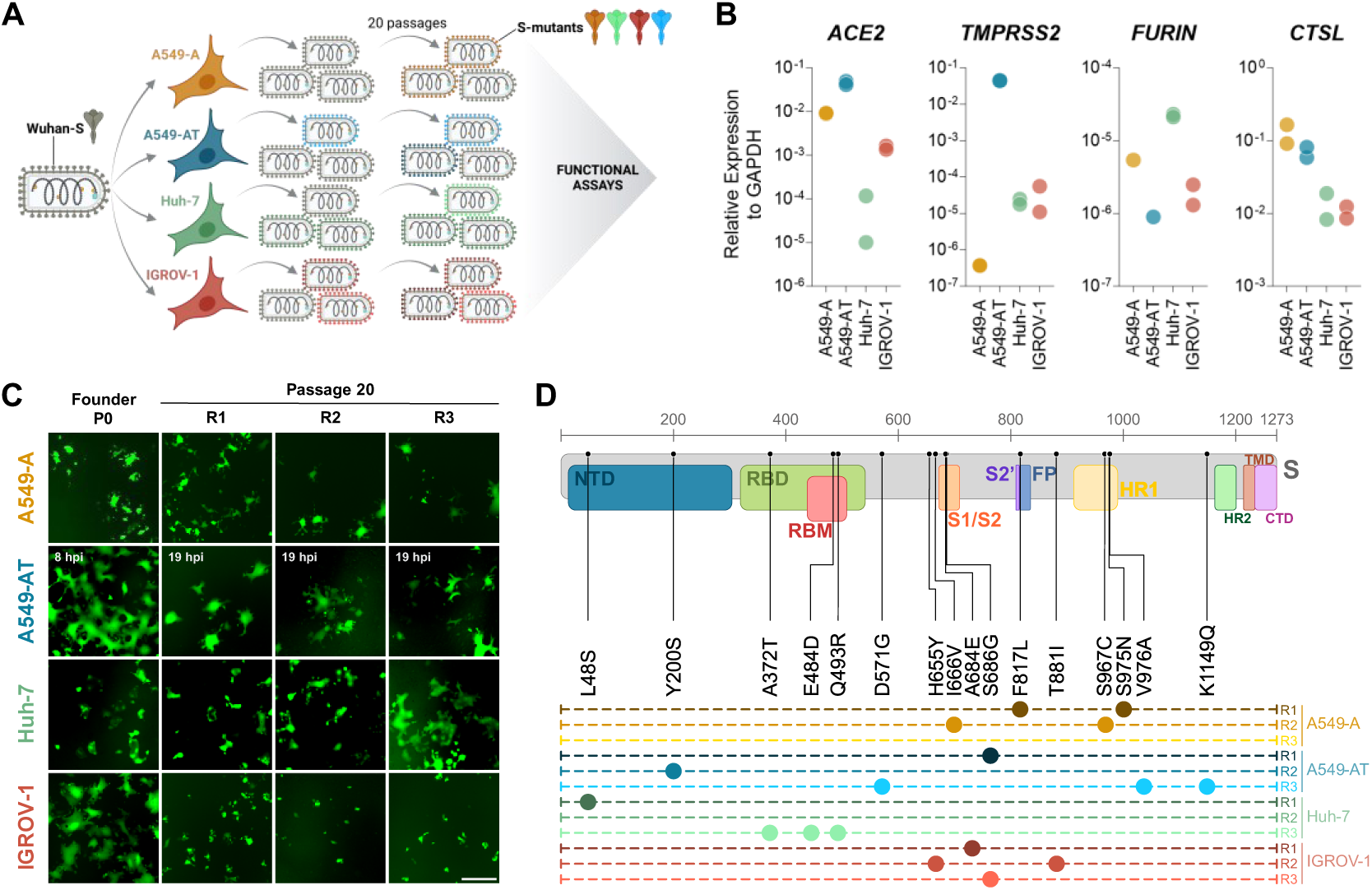
Experimental evolution of SARS-CoV-2 spike reveals cell type-specific adaptations. **(A)** Schematic representation of the experimental evolution. **(B)** Expression of *ACE2*, *TMPRSS2*, *FURIN* and *CTSL* mRNAs in the four cell lines used in the experimental evolution was measured by RT-qPCR. Each dot represents an independent experiment (*n* = 2). **(C)** Representative images of infection with the founder and evolved lineages (passage 20) in their own cell line. A549-A: 48 hpi, A549-AT: selected times indicated in each image, Huh-7: 16 hpi, IGROV-1: 16 hpi. Scale bar: 400 µm. **(D)** Mutations fixed in each lineage at passage 20 were identified by Sanger sequencing. Amino acid positions are indicated at the top in grey numbers and coloured boxes depict functional domains.

The experimental evolution consisted of 20 serial passages and was performed in triplicate for each cell line (R1-R3). In lineages evolved in A549-AT, Huh-7, and IGROV-1, we observed phenotypic changes in syncytia formation (**Figure 1C**). For example, the Huh-7_R3 evolved virus induced considerably more cell-cell fusion than the other two lineages evolved in Huh-7 or the founder virus. Conversely, all lineages evolved in IGROV-1 exhibited reduced cell-cell fusion and lineages evolved in A549-AT showed slower syncytia formation.

To identify mutations that were fixed during the evolution process, we Sanger-sequenced the S gene of the 12 evolved viruses (**Figure 1D**). In total, we found 16 amino acid substitutions, which mapped to different functional domains, including the N-terminal domain (NTD), the receptor binding domain (RBD), the S1/S2 junction, the S2’ cleavage region, and the heptad repeat 1 (HR1). Interestingly, we found substitutions H655Y and Q493R, which have been described in the Gamma and/or Omicron variants (**Table S1**). In contrast, the other mutations have not been reported to a significant frequency in nature^44^. Interestingly, though, one the changes (A372T) maps to a specific polymorphic site of the RBD suggested to play a key role in the origin of the SARS-CoV-2 pandemics^45^. Another mutation (E484D) involved a residue that has been strongly selected during SARS-CoV-2 adaptation to humans, but with different amino acid variants^46,47^. In total, we found three substitutions affecting the RBD (A372T, E484D and Q493R), all of which emerged in Huh-7 cells. We also found two mutations that disrupted the FCS (A684E and S686G), and two in its vicinity (H655Y and I666V). FCS-related mutations occurred at least once in all cell lines, except Huh-7. S686G arose independently in two lineages, strongly suggesting that FCS alteration was selectively advantageous in these cells.

### Spike mutations affect viral entry in a cell type-dependent manner

The above results revealed that the SARS-CoV-2 spike evolves distinctly in different cell lines, both at the phenotypic and genotypic levels. To explore this further, we used site-directed mutagenesis to recreate each individual mutation in the Wuhan-Hu-1 background, as well as five combinations of two or three mutations found in the evolved spikes. We then obtained VSV pseudotypes for each spike and titrated them in the four cell lines to assess the relative efficiency of viral entry (**Figure 2A**). This revealed that mutations affected entry in a cell type-dependent manner (mixed model effect with Geisser-Greenhouse correction: *p* < 0.0001). A striking example was the triple mutant D571G/V976A/K1149Q, which specifically increased viral entry in A549-AT, the cell line where this mutant emerged. Similarly, the entry of the triple mutant A372T/E484D/Q493R was particularly enhanced in Huh-7 cells, where this mutation was fixed. Finally, the H655Y/F881I was especially beneficial for viral entry in IGROV-1 cells, where it emerged. In all three cases, the corresponding single mutants affected viral entry to a lesser extent, suggesting epistasis between mutations. The S686G mutation appeared in IGROV-1 and A549-AT cells but increased pseudotype entry only in IGROV-1 cells. In A549-AT, this mutation may therefore confer a replicative advantage to the virus that is independent of viral entry.

**Figure 2.**
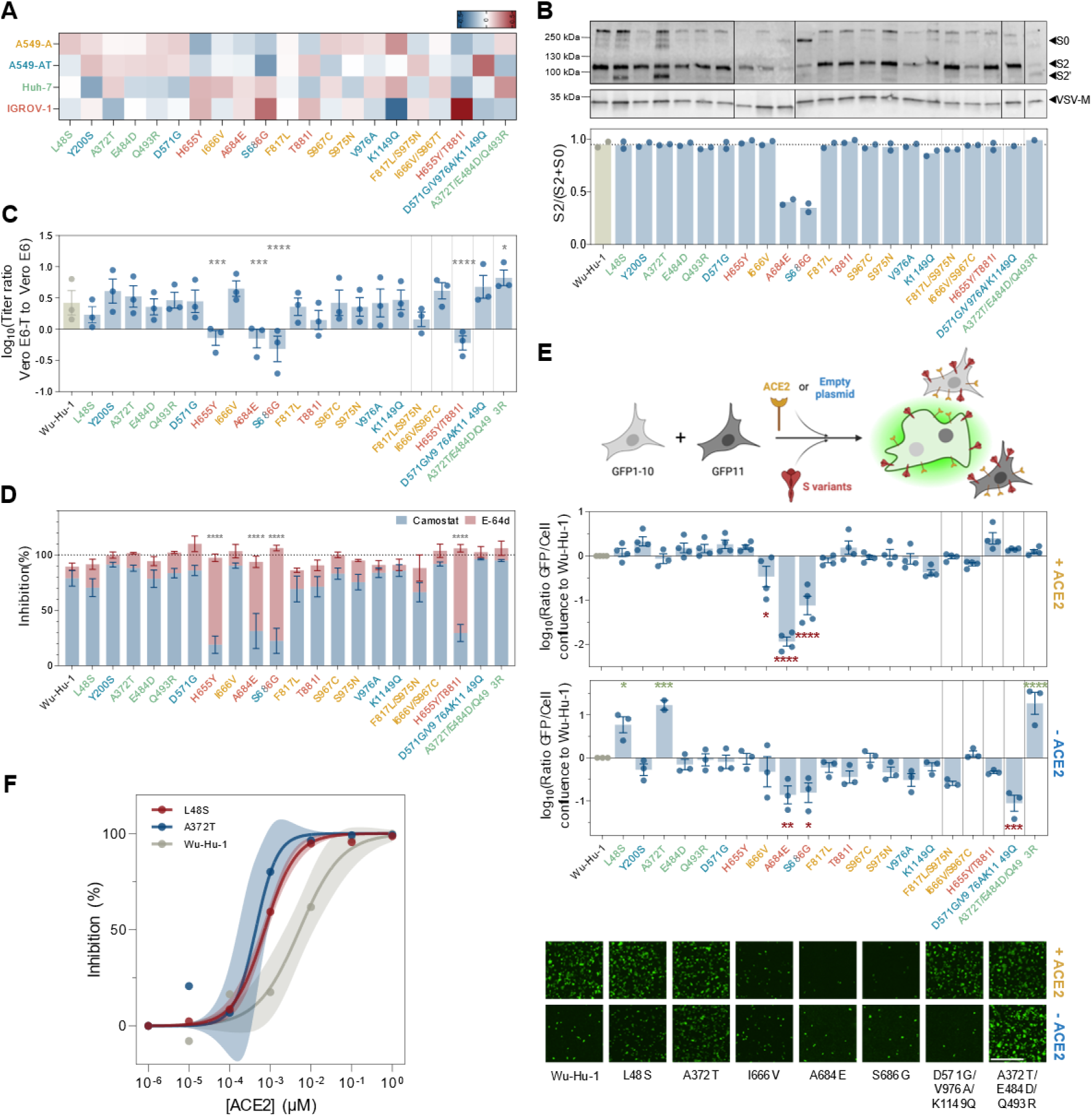
Functional characterization of mutations observed after 20 passages. **(A)** Heat map showing cell type dependent effects of each mutation on viral entry. VSV pseudotypes were titrated in the four cell lines. Within each cell line, titers were first normalized to the average titer of all mutants (log(titer) – mean(log(titer))). For each mutant, normalized titers were then normalized to their average across the four cell lines. Results represent the average of 2-3 independent experiments. Mutations are colored according to the cell line in which they were detected. **(B)** Anti-S2 Western Blot of pseudotyped VSV particles (top panel). Spike processing was calculated as the proportion of S2 over total spike (bottom panel). Each dot represents an independent blot (n=2). **(C)** TMPRSS2 usage of pseudotyped VSV particles was measured as the ratio of the titer in Vero E6-T to the titer in Vero E6 cells. Asterisks indicate the p-values obtained from a randomized-block one-way ANOVA with Dunnett’s correction for multiple comparisons, comparing all mutants to the Wuhan-Hu-1 reference. Data are shown as mean ± SEM, each dot corresponding to an independent experiment (n=3). **(D)** Infection inhibition reached with a 100 µM of camostat mesylate (blue) or E-64d (red) dose. Data are shown as mean ± SEM (n=3 independent experiments). Asterisks indicate the p-values obtained from a one-way ANOVA with Dunnett’s correction for multiple testing, comparing all mutants to the Wuhan-Hu-1 reference. **(E)** Cell-cell fusogenicity of the S mutants. A schematic of the GFP-Split assay is presented on top. The assay was performed in the presence (top graph) or absence (bottom graph) of human ACE2 overexpression. Data are shown as the log10 of the GFP confluence:cell confluence ratio normalized to that of the Wuhan-Hu-1 reference (mean ± SEM, n = 3-4 independent experiment). Asterisks show the p-values obtained from an ordinary one-way ANOVA with Dunnett’s multiple test correction, comparing all mutants to the Wuhan-Hu-1 reference. Representative images from statistically significant results are shown at the bottom. Scale bar: 800 µm. **(F)** Infection inhibition assay by soluble ACE2 against WT, L48S or A372T VSV pseudotypes. Each dot is the average of 3 technical replicates, lines correspond to a sigmoidal 2-parameter fit, and shaded areas correspond to 95% confidence intervals.

### S1/S2 mutations alter spike processing and entry pathway

To gain further insight into the functional consequences of these changes, we quantified the effect of all spike mutations on processing by furin, measured as the S2/(S2+S0) band ratio in Western Blots (**Figure 2B**). Consistent with previous reports^48^, we found that the two FCS mutations A684E and S686G decreased cleavage compared to the Wuhan-Hu-1 spike by approximately 50%.

In addition to S0 cleavage by furin, activation of the spike requires further processing at the S2’ site either by TMPRSS2 at the cell surface or by cathepsins in endosomes, depending on the entry route. Thus, we first assessed TMPRSS2 usage by comparing pseudotype titers in Vero E6 and Vero E6-T cells (**Figure 2C**). Whereas TMPRSS2 expression enhanced entry of the Wuhan-Hu-1 spike and most mutants by 1.4 - 4.8-fold, this effect was absent in the two FCS mutants, showing that TMPRSS2 activated these spikes less efficiently because they were not pre-processed by furin. A similar reduction in TMPRSS2 usage was observed for the H655Y substitution.

To confirm these results, we infected Vero E6-T cells with pseudotypes in the presence of the TMRPSS2 inhibitor camostat mesylate or the cathepsin inhibitor E-64d (**Figure 2D**). The Wuhan-Hu-1 spike and most mutants showed stronger inhibition by camostat than by E-64d, indicating that most S2’ cleavage was carried out by TMPRSS2. Conversely, the H655Y, A684E and S686G spikes were predominantly processed by cathepsins (78.5%, 62.5% and 83.8% inhibition by E-64d, respectively), in agreement with previous observations^49,50^.

These results confirm that the H655Y, A684E and S686G spikes enter the cell through endocytosis, whereas the Wuhan-Hu-1 spike and other variants predominantly fuse with the plasma membrane. By disrupting the FCS, substitutions A684E and S686G prevent cleavage by furin and hence reduce TMPRSS2 activity, restricting entry to the endosomal pathway. In contrast, the H655Y substitution does not prevent furin activity but nevertheless alters the viral entry route, probably through different mechanisms.

### Huh-7-acquired mutations increase cell-cell fusogenicity and binding to ACE2

In the above Western Blots (**Figure 2B**), the L48S and A372T spikes strikingly displayed an ∼ 80 kDa band, whose molecular weight is consistent with that of the S2’ cleavage fragment. This band was only slightly visible in pseudotypes bearing the Wuhan-Hu-1 spike or other mutants, suggesting that these two amino acid replacements promote spike processing. Interestingly, the L48S and A372T mutations were fixed in two independent lineages evolved in Huh-7 cells, which express the lowest levels of ACE2. We therefore hypothesized that these mutations might increase the activation of the SARS-CoV-2 spike or its affinity for ACE2.

To explore this, we first exploited the ability of the spike to induce cell-cell fusion upon interaction with ACE2, a process suggested to play a role in viral dissemination, immune escape and pathogenesis^7,51,52^. Specifically, we used a GFP complementation assay^53^ to measure the effect of each mutation on spike-mediated syncytia formation. In cells expressing high levels of ACE2 (HEK293T transfected with an ACE2 plasmid), most spikes induced cell-cell fusion to levels comparable to the Wuhan-Hu-1 spike. The only exception were FCS-disrupting mutations, which reduced syncytia formation by more than twofold (**Figure 2E**). This was expected, since these mutations promote entry through endocytosis, as shown above. In contrast, repeating these assays under very low levels of ACE2 (basal HEK293T expression^30,54^) revealed that the L48S and A372T spikes strongly increased cell-cell fusion compared to the Wuhan-Hu-1 spike (by 5.9- and 16.8-fold, respectively). The effect of A372T was particularly evident in the A372T/E484D/Q493R triple mutant (18.5-fold increase), which again supports epistasis between these mutations.

These results could be explained by an increased fusogenicity, but also by changes in receptor affinity, particularly for the RBD mutation A372T. As a surrogate test to measure binding affinity, we performed a competition assay with soluble human ACE2 (**Figure 2F**). Pseudotypes bearing Wuhan-Hu-1, L48S, or A372T spikes were incubated with increasing doses of soluble ACE2 prior to infection of Vero E6-T cells. Less soluble ACE2 was required to block the L48S (8.5-fold lower IC_50_) or A372T (15.4-fold lower IC_50_) spikes than the Wuhan-Hu-1. This shows that, in two independent replicates of evolution in Huh-7 cells, the SARS-CoV-2 spike became adapted to low ACE2 levels by acquiring mutations that increase interaction with the receptor and fusogenicity.

### Cell type-dependent effect of spike mutations on the fitness of recombinant viruses

To further test the cell type-dependent effects observed with pseudotypes, we assayed the effect of the most interesting spike mutations in the context of a replicative virus. For this, we constructed recombinant VSVs bearing the SARS-CoV-2 spike mutations L48S, A372T or S686G, which were selected for these analyses based on our above results. Whereas pseudotyped viruses allowed us to focus on entry, the fitness of these recombinant viruses could also depend on spike pre-processing during egress and syncytia induction.

Viral spread was monitored by measuring the area under the curve (AUC) of the GFP signal in infected cultures (**Figure 3A-B**), and viral titers were determined at different time points by the foci assay (**Figure 3C**). This showed that the L48S and A372T substitutions increased viral propagation relative to the Wuhan-Hu-1 control by 6.3- and 15.9-fold in Huh-7 cells, respectively (**Figure 3B**). Moreover, observation of the cell cultures showed that these two mutations increased cell-cell fusion in Huh-7 (**Figure 3A**), consistent with the results obtained above with low-ACE2 cells. In contrast, the L48S and A372T substitutions did not alter viral spread in other cell types. We also found that the two mutants replicated to higher titers than Wuhan-Hu-1 in Huh-7 (14.3- and 4.7-fold increase, respectively), whereas they were neutral in A549-A and A549-AT and even detrimental in IGROV-1 (**Figure 3C**). The latter is consistent with the strongly negative impact of A372T on viral entry in IGROV-1 (**Figure 2A**). Concerning the S686G substitution, it increased viral titers in A549-AT (27.3-fold) and IGROV-1 (10.4-fold) cells, the two cell types in which this mutation appeared (**Figure 3C**). The impact of this change on AUC values was negative in all cell types due to reduced cell-cell fusion (**Figure 2E**, **Figure 3A-B**), illustrating that viral spread and progeny production measures are not necessarily coupled. The S686G mutation did not increase viral entry in A549-AT (**Figure 2A**), suggesting that its beneficial effect on viral fitness is likely related to its lower cell-cell fusogenicity that may be beneficial in this fast-fusing cell line.

**Figure 3.**
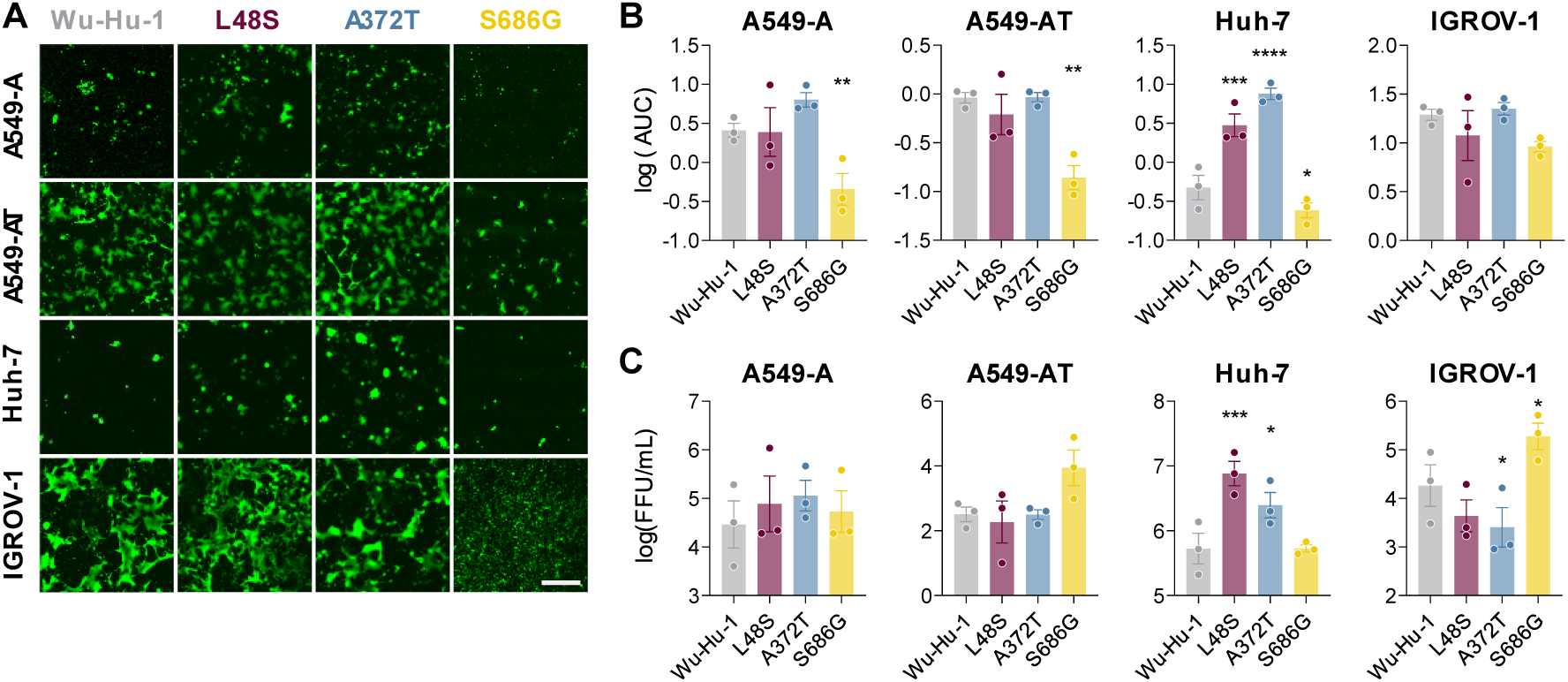
Cell type-dependent effect of spike mutations on the fitness of recombinant viruses. **(A)** Representative images of infection of the four cell lines with VSVs expressing Wuhan-Hu-1 or mutant spikes. Images were acquired at different time points for each cell line (92 hpi in A549-A, 9 hpi in A549-AT, 48 hpi in IGROV-1 and 76 hpi in Huh-7). Scale bar: 800 µm. **(B)** Viral spread in the cell culture was quantified by measuring AUC over the course of infection. Data are mean ± SEM are shown, each dot corresponding to an independent experiment (n=3). Asterisks correspond to p-values obtained from a randomized-block one-way ANOVA with Dunnett’s correction for multiple tests, comparing all mutants to the Wuhan-Hu-1 reference. **(C)** Titration of supernatants at 12 hpi (A549-AT), 48 hpi (IGROV-1) or 72 hpi (A549-A and Huh-7). Data are shown as mean ± SEM, each dot corresponding to an independent experiment (n=3). Asterisks correspond to p-values obtained from a randomized-block one-way ANOVA with Dunnett’s correction for multiple tests, comparing all mutants to the Wuhan-Hu-1 reference.

## Discussion

We have experimentally evolved the SARS-CoV-2 spike protein in four different human cell lines to identify cell type-specific adaptation mechanisms. Of the 16 mutations that became fixed after 20 passages, some had a clear effect on entry efficiency, spike processing, entry pathway usage, cell-cell fusion, or ACE2 affinity. However, many mutations had no effect in any of these functional assays. These mutations could have evolved through genetic hitch-hiking, or their fitness benefits could be unrelated to viral entry, or not revealed by the assays performed. For example, we did not measure the ACE2 affinity of all mutants because it was prohibitive to produce a large number of soluble spikes to measure affinity through standard methods (e.g., ELISA, SPR or BLI). Similarly, other processes that may play a role in spike adaptation were not evaluated, such as binding to attachment factors other than ACE2, intracellular spike trafficking, and glycosylation.

Our evolution experiment in cell cultures recovered two mutations known to be important for viral fitness in nature. H655Y and Q493R have been found in globally circulating variants of concern, such as Gamma and Omicron. It is interesting to note that the Q493R mutation, which has been associated with antibody escape^55^, evolved here in the absence of immune pressure, suggesting that antibody escape was probably not a major factor driving the evolution of this amino acid substitution. Indeed, Q493R has also been shown to increase binding to mouse ACE2^56,57^, highlighting that a single spike mutation can have multiple effects. The H655Y mutation, which emerged in IGROV-1 cells, has been found to stabilize the spike on virions and to shift the RBD towards the open conformation required for ACE2 binding^58^. Interestingly, IGROV-1 cells are particularly susceptible to the Omicron variant^43^, consistent with our finding that the H655Y mutation evolved only in these cells. Similar effects have been reported for other well-known mutations, such as D614G, which increases SARS-CoV-2 infectivity and transmissibility by stabilizing S1/S2 dimers on virions^59,60^. The D614G substitution was fixed very early after the introduction of SARS-CoV-2 in humans and has been maintained in all SARS-CoV-2 B lineages. However, we did not find this change in our experimentally evolved spikes, as well as many other important mutations reported in vivo. The stochastic nature of evolution, the small number of infection cycles elapsed in our experiments, or differences between selective pressures in vivo and in cell cultures can explain these discrepancies. Moreover, although the spike-encoding recombinant VSV used here mimics SARS-CoV-2 entry^34^, the spike C-terminal deletion, the VSV genome, and the fact that VSV buds at the plasma membrane whereas SARS-CoV-2 likely buds at the ERGIC may impact spike evolution. Although most of the mutations observed here have not yet been identified at high frequency in humans, their functional characterisation will help to understand their role if they emerge in future variants.

SARS-CoV-2 is the only sarbecovirus that possesses a multibasic FCS in the S1/S2 region. The FCS has been shown to be essential for efficient transmissibility, which may explain why it has been maintained throughout SARS-CoV-2 evolution^61^. However, in cell cultures, the FCS has been repeatedly shown to be inactivated upon SARS-CoV-2 passaging^62,63^. Of the 16 different mutations fixed in our evolved viruses, two directly disrupted the FCS and one (S686G) occurred in two independent lineages. The functional effect of FCS mutations (e.g., S686G) was cell type-dependent, in agreement with previous reports^64^. An obvious effect of FCS loss is that TMPRSS2 processing is no longer an efficient spike activation mechanism, and that entry through the endocytic route is enforced. In cell cultures, a possible advantage of this shift is avoidance of syncytia formation, since extensive cell-cell fusion may compromise cell viability and viral progeny production^7,65^. This advantageous effect might also take place to some extent in vivo. For example, although we did not see an effect of the single H655Y mutation on cell-cell fusion, it has been demonstrated that in the context of the Omicron spike, this mutation was responsible for decreased fusogenicity^50^. We thus speculate that the Omicron/Gamma mutation H665Y might be an evolutionary solution to enable preferential use of the endocytic route and decrease excessive syncytia formation without disrupting the FCS.

Mutations affecting the RBD (A372T, E484D and Q493R) appeared only in Huh-7 cells, which expressed the lowest levels of ACE2 of the four cell lines. This suggests that the spike-ACE2 interaction is not a major evolutionary pressure in the other cell types but that in Huh-7, the virus may have evolved either to maximize interaction with ACE2 or to avoid dependency on ACE2 for viral entry. Whereas E484D has not been reported at a significant rate in nature, other mutations at this residue have been fixed several times. For example, E484K arose in the Beta and Gamma variants and E484A was fixed in all Omicron subvariants. Both mutations have been linked to immune escape^46^. E484D has nevertheless been previously reported in cell cultures, and suggested to allow ACE2-independent viral entry^39,66–68^. Recent work has shown that E484D strongly increases interaction with TMEM106B, an alternative SARS-CoV-2 receptor that mediates ACE2-independent infection^21,69^. Our results therefore suggest that, in the presence of low levels of ACE2 in Huh-7, the SARS-CoV-2 spike acquired the E484D mutation to favor interactions with TMEM106B or other unidentified alternative receptors. Since these effects could in principle expand the SARS-CoV-2 tropism, it is surprising that this mutation has not been reported in nature.

Another RBD mutation acquired in Huh-7 is A372T. All SARS-CoV-2-related viruses have a threonine at position 372 of their spike protein. Kang et al. proposed that a T372A change occurred early in the viral cross-species transmission event and was rapidly favored by natural selection^45^. This previous work showed that the A372T mutation decreased viral replication and binding affinity to ACE2. However, our results indicate that A372T increases interaction with ACE2, leading to increased cell-cell fusion. These discrepancies are probably due to technical differences. While Kang et al. assessed the RBD by ELISA, we used a soluble ACE2 competition assay with viral pseudotypes containing entire spikes. Our results therefore suggest that in the context of a functional spike, the threonine variant increases interaction with ACE2, either by enhancing binding affinity or by stabilizing S1/S2 dimers. The latter is supported by recent data showing that the introduction of a threonine at this site in the SARS-CoV-2-related BANAL-20-52 spike decreases viral shedding^70^. Finally, the fitness advantage of the A372T mutant was highly specific to Huh-7. Interestingly, Huh-7 was also the cell line expressing the highest levels of furin. We thus speculate that, in viruses egressed from these cells, the spike is strongly processed by furin and that the A372T stabilizes S1/S2 dimers, preventing S1 shedding. The L48S mutation also occurred in Huh-7 and had a similar effect on cell fusion, ACE2 binding and viral fitness. It would therefore be interesting to study how this substitution affects S1/S2 stability.

In conclusion, our data show that SARS-CoV-2 spike adaptation is cell type-dependent, illustrating how replication in different tissues may contribute to viral diversification. Further work aimed at elucidating the interplay between viral and host factors in different cellular environments will be crucial for understanding the complex dynamics of SARS-CoV-2 evolution.

## Methods

### Cells

A549-ACE2 (A549-A), Huh-7, A549-ACE2-TMPRSS2 (A549-AT; Invivogen a549-hace2tpsa), Vero E6 (kind gift of Dr. Luis Enjuanes, Centro Nacional de Biotecnología), Vero E6-TMPRSS2 (Vero E6-T; JCRB Cell Bank JCRB1819), HEK-293T-GFP1-10 and HEK-293T-GFP-11 (a gift from Olivier Schwartz, Institut Pasteur), BHK-21 and BHK-G43 (BHK21 cells that can be induced to express the VSV G protein^71^; kind gift of Dr. Gert Zimmer) were grown in DMEM (Gibco™ 10502703) supplemented with 10% fetal bovine serum (FBS, Gibco™ 10270106), 1% non-essential amino acids (Gibco™ 11140050), penicillin and streptomycin (10 U/mL and 10 µg/mL, respectively; Gibco™ 15140122), and amphotericin B (250 ng/mL, Gibco™ 15290026). IGROV-1 were grown in RPMI 1640 medium (Gibco™ 21875091) supplemented with 10% FBS, penicillin (10 U/mL), streptomycin (10 µg/mL), and amphotericin B (250 ng/mL). Additionally, the following antibiotics were used: 0.5 mg/mL geneticin (Gibco™ 10131027) in Vero E6-T, 0.1 µg/mL blasticidin S (Gibco™ R21001) in A549-A, 0.5 µg/mL puromycin (Gibco™ A1113803) and 0.3 mg/mL hygromycin B (Gibco™ 10687010) in A549-AT, 1 µg/mL puromycin in HEK-293T-GFP1-10 and HEK-293T-GFP-11, and of 500 µg/mL hygromycin B and 1 mg/mL zeocin in BHK-G43. In all cases, cells were maintained at 37°C and 5% CO_2_ in a humidified incubator and were regularly shown to be free of mycoplasma contamination by PCR.

### Gene expression analysis

For gene expression analysis, total RNA was harvested from confluent p100 dishes. Briefly, 2.5 mL RNAzol was added and incubated for 10 min at room temperature, followed by centrifugation at 12,000 g for 15 min. One volume of 100% 2-propanol was added to the supernatant, incubated for 10 min at room temperature and centrifuged at 12,000 g for 15 min. Supernatants were discarded and pellets were washed twice by adding 1 mL 75% ethanol and centrifuging at 6,000 g for 2 min. Finally, RNA was solubilized with 10 µL RNAse-free water. For qPCR assays, 100 ng RNA were used as template for the TaqPath 1-Step RT-qPCR assay (Applied Biosystems, A15299), following manufacturer’s instructions and specific, commercial ready-to-use primers and probes for each *ACE2*, *TMPRSS2*, *FURIN*, *CTSL*, and *GAPDH* from IDT PrimeTime™ Predesigned qPCR Assays (Hs.PT.58.27645939; Hs.PT.58.39738666; Hs.PT.58.1294962; Hs.PT.58.39748218; Hs.PT.39a.22214836). Results were normalized to GAPDH expression (Δ*C_t_* = *C_t_*_gene_ - *C_t_*_GAPDH_) and expressed as 2^-Δ*Ct*^.

### Recombinant viruses

Recovery of replication-competent recombinant VSV bearing Wuhan-S (wild-type or mutants) was performed as follows. Briefly, BHK-G43 cells (BHK-21 cells that express the VSV glycoprotein after mifepristone treatment^71^) were seeded at a density of 1.5·10^5^ cells/mL in DMEM 5% FBS without antibiotics in 12-well plates (1 mL per well). The following day, the viral genome pVSVeGFP-ΔG-Wu-S-ΔCt (Wuhan-Hu-1 with a deletion of the last 21 amino acids; 25 fmol) was co-transfected with helper plasmids encoding VSV P (25 fmol), N (75 fmol) and L (25 fmol) proteins and the T7 RNA polymerase (50 fmol) using lipofectamine 3000 (Invitrogen™ L3000150) for 3 h at 37°C. Then the medium was replaced with DMEM 10% FBS supplemented with 10 nM mifepristone to induce VSV-G expression and cells were incubated at 33°C for 36 h, followed by 48 h at 37°C. Supernatants from GFP-positive cells were harvested, clarified by centrifugation at 10,000 g for 10 min, and used to infect a fresh VSV-G-induced BHK-G43 p100 dish culture for amplification. Supernatants were harvested at 24-48 hours post-infection (hpi) and clarified in the same way. To clean viruses from VSV-G, non-G expressing cells (BHK-21 or Vero E6) were inoculated with the BHK-G43-amplified viruses, inoculum was removed, cells were washed 5 times with PBS and incubated in DMEM containing 2% FBS and 25% of an anti-VSV-G neutralizing monoclonal antibody obtained in-house from a mouse hybridoma cell line, as described previously^72^. Supernatants were harvested at 24-48 hpi, clarified by centrifugation, and stored at −80°C.

### Experimental evolution

Experimental evolution was started from the Wuhan-Hu-1-ΔCt Spike-expressing recombinant VSV and was performed in independent triplicates in A549-A, A549-AT, Huh-7, and IGROV-1. Six-well plates confluent monolayers were infected with 300 µL of the virus at an MOI < 0.01. After 48 h, supernatants were harvested, used to infect fresh 6-well cultures and titrated, until 20 serial passages were completed. For titration of each evolution passage, serial dilutions were used to infect 50,000 Vero E6-T cells in suspension in a total volume of 200 µl in 96-well plates. After 24 h, plates were imaged in the Incucyte SX5 Live-Cell Analysis System (Sartorious), GFP foci were counted, and viral titers were expressed as FFU/mL.

### Sequencing

Evolved lineages were Sanger-sequenced at passage 20. For that, 150 µL of infection supernatant were harvested and clarified by centrifugation at 2,000 g for 10 min. RNA was harvested using the NucleoSpin RNA Virus kit (MACHEREY-NAGEL, 740956) following manufacturer’s instructions and reverse-transcribed with SuperScript IV (Invitrogen) using a specific primer (5’-CTCGAACAACTAATATCCTGTC-3’). cDNA (10 ng) was used for amplification of the spike gene with Phusion hot Start II High-Fidelity PCR Master Mix (ThermoFisher) using primers 5’-CTCGAACAACTAATATCCTGTC-3’ and 5’-GTTCTTACTATCCCACATCGAG-3’. Amplification product was Sanger-sequenced using PCR primers, a VSV backbone primer 5’-GGTCTCGAGCGTGATATCTG-3’ and S primers 5’-TTGCTGTATGACCAGTTGCTG-3’, 5’-GAAAGTACTACTACTCTGTATGGTTGG-3’, 5’-GGTTTAATTGTGTACAAAAACTGCC-3’, and 5’-TTCTGCAGCTCTAATTAATTGTTGA-3’.

### Viral growth assays

IGROV-1, Huh-7, A549-A, and A549-AT were plated in 12-well plates at 50% confluence. The following day, cells were infected with 100 µL of virus at 0.01 multiplicity of infection (MOI). Plates were imaged every 3 h for the first 24 hpi and every 6 h afterward in an Incucyte SX5 Live-Cell Analysis System (Sartorious). GFP area and cell confluence data were obtained with the Incucyte analysis software and the infected cells ratio was calculated as the ratio of both. The area under the curve (AUC) was calculated between *t*_0_ and 82 - 94 hpi (A549-A), 9 - 14 hpi (A549-AT), 62 - 76 hpi (Huh-7) or 48 - 64 hpi (IGROV-1). Time points were adjusted between experimental replicates to account for variability in infection kinetics. The same time point was used to compare all mutants. Input virus (*t*_0_) and supernatants from *t*_12h_ (A549-AT), *t*_48h_ (IGROV-1) and *t*_72h_ (Huh-7 and A549-A) were titrated in Vero E6-TMPRSS2 in 24-well plates. Briefly, 100 µL of serial dilutions were used to infect confluent monolayers. After 1 h of incubation at 37°C, cells were overlaid with DMEM 2% FBS with 0.5% agar. Plates were imaged 24 h in an Incucyte SX5 Live-Cell Analysis System (Sartorious), GFP counts were quantified using the Incucyte analysis software and titers were expressed as focus forming units per mL (FFU/mL).

### Site-directed mutagenesis

Site-directed mutagenesis of SARS-CoV-2 spike was performed with the QuikChange II XL Site-Directed Mutagenesis Kit (Agilent 200522), using primers listed in **Supplementary table S2** and according to manufacturer’s instructions. For each reaction, 10 ng of template was mutated using 250 ng of each primer pair (125 ng each) and 2.5 U/µL of PfuUltra HF DNA polymerase, and the PCR cycling parameters as recommended. PCR products were digested at 37°C for 1 h with *Dpn*I (Thermo Scientific™, FD1704) and transformed into NZY5α competent cells (Nzytech MB00402). The presence of the mutation and the absence of undesired mutations were confirmed by Sanger sequencing.

### VSV pseudotyping

Approximately 8·10^6^ HEK-293T cells were seeded into T75 flasks previously coated with poly-D-lysine (Gibco™ A3890401). After 16 - 24 h (when cells reached around 90% confluence), cells were transfected with 30 µg of His-tagged, eGFP-encoding, codon-optimized spike expression plasmid using Lipofectamine 2000 (Invitrogen™ 11668019) following manufacturer’s instructions. At 24 h post-transfection, cells were infected with a VSVΔG+G virus at MOI = 3. Supernatants containing pseudotypes were harvested at 24 hpi, cleared by centrifugation at 2,000 g for 10 min, passed through a 0.45 µm filter, aliquoted and stored at −80°C.

### Pseudotype titration

Pseudotypes were serially diluted in 1x DMEM supplemented with 2% FBS, preincubated 1:1 with an anti-VSV-G neutralizing monoclonal antibody for 30 min at 37 °C and used to infect confluent monolayers of A549-A, A549-AT, Huh-7, IGROV-1, Vero E6 or Vero E6-T cells. Plates were imaged at 24 hpi using the Incucyte SX5 Live-Cell Analysis System (Sartorious), GFP-positive cells were automatically detected using the Incucyte analysis software, and titers were expressed as infectious units per mL (IU/mL). For TMPRSS2 usage analysis, the ratio between Vero E6-T and Vero E6 titers was calculated.

To study cell type-specific effects of each mutation on viral entry, titers were first normalized within each cell line to the mean titers of all mutants (log(titer) – mean(log(titer))), and then a second normalization was performed for each mutant to its own average across the four cell lines. This normalization controls for absolute titers and possible differences in spike incorporation and focuses on the relative capacity of the spike protein to mediate viral entry in different cell lines.

### Western blot

One mL volume of pseudotyped or recombinant virus-containing supernatant was pelleted by high-speed centrifugation (30,000 g for 2h) and lysed in 30 µL of lysis buffer (cell lysis buffer II (Invitrogen™ FNN0021) supplemented with cOmplete protease inhibitor (Roche 11836170001)) for 30 min on ice. Approximately 10^6^ cells were washed with cold PBS and lysed in 80 µL of lysis buffer for 30 min on ice. Cell lysates were cleared by centrifugation at 15,000 g for 10 min at 4°C. Lysates were diluted in 4X Laemlli buffer (Bio-Rad) supplemented with 10% mercaptoethanol and denatured at 95°C for 5 min. Denatured samples (15 µL) were separated by SDS-PAGE using pre-cast 4 - 20% gels (Mini.PROTEAN TGX Gels, Bio-Rad) and transferred onto a 0.45 µm PVDF membrane (Thermo Scientific™). Membranes were blocked for 1 h at room temperature (r.t.) or overnight at 4°C in 3% BSA TBS-T (20 mM tris, 150 mM NaCl, 0.1% Tween-20, pH 7.5), and then incubated for 1 h at r.t. with the following antibodies: mouse anti-6xHis (1:1,000, MA1-21315, Invitrogen), mouse anti-VSV-M (1:1,000, [23H12], Kerafast), or rabbit anti-GAPDH (1:2000, ABS16, Merck Millipore). Primary antibodies were detected using goat Cy3-conjugated anti-rabbit IgG (1:10,000, A10520, Thermofisher) or goat Cy5-conjugated anti-mouse-IgG (1:10,000, A10524, ThermoFisher). Images were acquired on an AMERSHAM ImageQuant 800 (GE Healthcare) and analyzed with the Image Studio Lite software (LICOR, v5.2).

### Selective inhibition of alternative entry pathways

Vero E6-T cells were plated in a 96-well plate. The following day, cells were pre-treated with 100 µM of camostat mesylate (TMPRSS2 inhibitor; Sigma-Aldrich SML0057) or E-64d (cathepsin inhibitor; Sigma-Aldrich E8640) for 1 h at 37°C. Cells were then infected in duplicate with 1,000 IU of pseudotype (pre-incubated with anti-VSV-G antibody for 30 min at 37°C). Plates were scored for GFP foci 24 hpi, areas were quantified using the Incucyte analysis software, and infection inhibition was calculated relative to a mock-treated control. The percentage of inhibition was calculated as 100x (GFP area with mock - GFP area with inhibitor)/(GFP area with mock).

### Cell-cell fusion

The cell-cell fusion assay was performed as previously described^53^. Briefly, HEK-293T-GFP1-10 and HEK-293T-GFP-11 were mixed at a 1:1 ratio (6·10^5^ cells per well of a 96-well plate) and were co-transfected with 50 ng of S (WT or mutant) plasmid, and 50 ng of human ACE2-encoding plasmid or an empty plasmid control using Lipofectamine 2000 (Invitrogen) following manufacturer’s instructions. Plates were placed at 37°C and 5% CO_2_ in an Incucyte SX5 Live-Cell Analysis System (Sartorious). GFP signal and phase contrast images were analysed at 18-36 hpi. The percentage of fusion was calculated as the ratio between GFP area and cell confluence measured using the Incucyte analysis software.

### Infection inhibition by soluble hACE2

The soluble peptidase domain of human ACE2 (residues 19-615) was produced in a system of baculovirus/insect cells and purified at the IBV-Covid19-Pipeline (Institute of Biomedicine of Valencia-CSIC) as previously reported^73^. Purified protein was concentrated by centrifugal ultrafiltration using Amicon Ultra 30 K (Millipore) and quantified spectrophotometrically at 280 nm (mass extinction coefficient E 1%: 21.9 g-1 L cm-1). Ca. 5000 IU of each pseudotype were pre-incubated 1:1 with anti-VSV-G antibody and with serial dilutions of the soluble human ACE2 or a vehicle (PBS) control for 1 h at 37°C, and then used to infect Vero E6-T cells seeded in 96-well plates. The quantity of GFP-positive cells at 24 hpi was measured using the Incucyte SX5 Live-Cell Analysis System (Sartorious) and the Incucyte analysis software. The percentage of infection inhibition, an indirect measure of binding affinity to ACE2, was calculated as 100x (GFP-positive cells with vehicle – GFP-positive cells with hACE2)/(GFP-positive cells with vehicle).

### Statistics

All statistical tests were performed using GraphPad Prism 9.0 software. Specific tests are specified in figure legends. A *p*-value threshold for statistical significance of 0.05 was used in all cases. For all statistical tests, significance thresholds were: ns, not significant; *, *p* < 0.05; **, *p* < 0.01; ***, *p* < 0.001; ****, *p* < 0.0001.

## Author contributions

J.D. and R.S. designed research; M.C.-S. performed research; M.-C.M., R.G., S.F.E. and A.M. provided critical reagents and methodology, M.C.-S., J.D., and R.S. analyzed data; M.C.-S., J.D. and R.S. wrote the paper; A.R. and R.S. provided funding.

## Acknowledgments

We thank Carlos Baeza-Delgado for technical assistance. This work was financially supported by Advanced Grant 101019724 - EVADER from the European Research Council (ERC) and grant PID2020-118602RB-I00 – ZooVir from the Spanish Ministerio de Ciencia e Innovación to R.S. M.C.-S. was funded by a Ph.D. fellowship from the Spanish Ministerio de Ciencia e Innovación, the Agencia Estatal de Investigación and the European Social Fund Plus (PRE2021-099824). J.D. was funded by an EMBO postdoctoral fellowship (ALTF 140-2021). R.G. acknowledges funding by grants from the European Commission NextGenerationEU fund (EU 2020/2094), through CSIC’s Global Health Platform (PTI Salud Global). A.R. and R.S. received funding from the Fundació La Marató de TV3 (202130-31).

## Consortia

### The IBV-Covid19-Pipeline

Anmol Adhav, Clara Marco-Marin, Laura Villamayor-Bellichón, Carolina Espinosa, Maria del Pilar Hernández-Sierra, Rafael Ruiz-Partida, Nadine Gougeard, Alicia Forcada-Nadal, Sara Zamora-Caballero, Antonio Rubio-del-Campo, Roberto Gozalbo-Rovira, Carla Sanz-Frasquet, Francisca Gallego del Sol, Alonso Felipe-Ruiz, María Luisa López-Redondo, Santiago Ramón-Maiques, Jeronimo Bravo, Vicente Rubio, José Luis Llácer, Alberto Marina

