## Supplementary information for "Cell type-specific adaptation of the SARS-CoV-2 spike"

### Supplementary Materials

Tables S1 to S2.

### Supplementary Materials

**Table S1.** Mutations detected by Sanger sequencing in the 12 evolved lineages (passage 20).

| Base substitution | Amino acid subtitution | Cell line | Nextstrain count [a] | Associated VOC |
|---|---|---|---|---|
| T143C | L48S | Huh-7 | 4 | |
| A599C | Y200S | A549-AT | 7 | |
| G1114A | A372T | Huh-7 | 31 | |
| A1452T | E484D | Huh-7 | 169 | |
| A1478T | Q493R | Huh-7 | 1410013 | Omicron (BA.1, BA.5) |
| A1712G | D571G | A549-AT | 26 | |
| C1963T | H655Y | IGROV-1 | 1568908 | Gamma, Omicron |
| A1996G | I666V | A549-A | 156 | |
| C2051A | A684E | IGROV-1 | 8 | |
| A2056G | S686G | A549-AT, IGROV-1 | 31 | |
| T2449C | F817L | A549-A | 102 | |
| C2642T | T881I | IGROV-1 | 26 | |
| A2899T | S967C | A549-A | 1 | |
| G2924A | S975N | A549-A | 20 | |
| T2927C | V976A | A549-AT | 16 | |
| A3445C | K1149Q | A549-AT | 19 | |

[a]accessed on July 19, 2023

**Table S2.** Primers used for site directed mutagenesis. In yellow, specific changes introduced.

| | Primer | Sequence (5'-3') |
|---|---|---|
| Pseudoviruses | L48S-Fw | GTTCAGATCCAGCGTGTCTCACTCTACCCAGGACC |
| | L48S-Rv | GGTCCTGGGTAGAGTGAGACACGCTGGATCTGAAC |
| | Y200S-Fw | GAACATCGACGGCTCCTTCAAGATCTACAGCAAGC |
| | Y200S-Rv | GCTTGCTGTAGATCTTGAAGGAGCCGTCGATGTTC |
| | A372T-Fw | GTGCTGTACAACTCCACCAGCTTCAGCACCTTCAAG |
| | A372T-Rv | CTTGAAGGTGCTGAAGCTGGTGGAGTTGTACAGCAC |
| | E484D-Fw | CTTGTAACGGCGTGGATGGCTTCAACTGCTAC |
| | E484D-Rv | GTAGCAGTTGAAGCCATCCACGCCGTTACAAG |
| | Q493R-Fw | GCTACTTCCCACTGAGGTCCTACGGCTTTCAGCCC |
| | Q493R-Rv | GGGCTGAAAGCCGTAGGACCTCAGTGGGAAGTAGC |
| | D571G-Fw | GGCCGGGATATCGCCGGCACCACAGACGCCGTTAG |
| | D571G-Rv | CTAACGGCGTCTGTGGTGCCGGCGATATCCCGGCC |
| | T632A-Fw | GATCAGCTGACACCTGCATGGCGGGTGTACTCCAC |
| | T632A-Rv | GTGGAGTACACCCGCCATGCAGGTGTCAGCTGATC |
| | H655Y-Fw | GTCTGATCGGAGCCGAGTACGTGAACAATAGCTACG |
| | H655Y-Rv | CGTAGCTATTGTTCACGTACTCGGCTCCGATCAGAC |
| | I666V-Fw | CGAGTGCGACATCCCCGTGGGCGCTGGCATCTGTG |
| | I666V-Rv | CACAGATGCCAGCGCCCACGGGGATGTCGCACTCG |
| | A684E-Fw | CAAACAGCCCCAGACGGGAGAGATCTGTGGCCAGC |
| | A684E-Rv | GCTGGCCACAGATCTCTCCCGTCTGGGGCTGTTTG |
| | S686G-Fw | GCCCCAGACGGGCCAGAGGCGTGGCCAGCCAGAGC |
| | S686G-Rv | GCTCTGGCTGGCCACGCCTCTGGCCCGTCTGGGGC |
| | F817L-Fw | GCCCAGCAAGCGGAGCCTGATCGAGGACCTGCTG |
| | F817L-Rv | CAGCAGGTCCTCGATCAGGCTCCGCTTGCTGGGC |
| | F881I-Fw | GCCCTGCTGGCCGGCATCATCACAAGCGGCTGGAC |
| | F881I-Rv | GTCCAGCCGCTTGTGATGATGCCGGCCAGCAGGGC |
| | S967T-Fw | CCCTGGTCAAGCAGCTGACCTCCAACTTCGGCGCC |
| | S967T-Rv | GGCGCCGAAGTTGGAGGTCAGCTGCTTGACCAGGG |
| | S975N-Fw | CTTCGGCGCCATCAGCAACGTGCTGAACGATATCC |
| | S975N-Rv | GGATATCGTTCAGCACGTTGCTGATGGCGCCGAAG |
| | K1149Q-Fw | CCGAGCTGGACAGCTTCCAGGAGGAACTGGATAAG |
| | K1149Q-Rv | CTTATCCAGTTCCTCCTGGAAGCTGTCCAGCTCGG |
| Recombinant VSV | L48S-Fw | CAGATCCTCAGTTTCACATTCAACTCAGGACTTG |
| | L48S-Rv | CAAGTCCTGAGTTGAATGTGAAACTGAGGATCTG |
| | A372T-Fw | CTGTCCTATATAATTCCACATCATTTTCCAC |
| | A372T-Rv | GTGGAAAATGATGTGGAATTATATAGGACAG |
| | S686G-Fw | CTCGGCGGGCACGTGGTGTAGCTAGTC |
| | S686G-Rv | GACTAGCTACACCACGTGCCCGCCGAG |
